# Single cell sequencing data identify distinct B cell and fibroblast populations in stricturing Crohn’s disease

**DOI:** 10.1101/2023.09.04.556163

**Authors:** David T Humphreys, Amy Lewis, Belen Pan-Castillo, Giulio Berti, Charles Mein, Eva Wozniak, Hannah Gordon, Radha Gadhok, Annamaria Minicozzi, Joanna ChinAleong, Roger Feakins, Eleni Giannoulatou, Louisa K James, Andy J Stagg, James Oliver Lindsay, Andrew Silver

**Affiliations:** Victor Chang Cardiac Research Institute, Sydney, NSW 2010, Australia; St Vincent’s Clinical School, University of New South Wales, Sydney, NSW 2052, Australia; Centre for Genomics and Child Health, Blizard Institute, Barts and The London School of Medicine & Dentistry, London E1 2AT UK; Genome Centre, Blizard Institute, Barts and The London School of Medicine & Dentistry, London E1 2AT UK; Centre for Immunobiology, Blizard Institute, Barts and The London School of Medicine & Dentistry, London E1 2AT UK; Department of Colorectal Surgery, Division of Surgery & Perioperative Care, The Royal London Hospital, Whitechapel, London E1 1BB; Department of Histopathology, The Royal London Hospital E1 1BB UK; Department of Cellular Pathology, Royal Free London NHS Foundation Trust, London NW3 2QG, UK

## Abstract

We used human full thickness Crohn’s disease (CD) small bowel resection specimens and single cell RNA sequencing to identify potential therapeutic targets for stricturing (S)CD. Using an unbiased approach, 16 cell lineages were assigned within 14,539 sequenced cells from patient-matched SCD and non-stricturing (NSCD) preparations. SCD and NSCD contained identical cell types. Amongst immune cells, B cells and plasma cells were selectively increased in SCD samples. B cell subsets suggested formation of tertiary lymphoid tissue in SCD and compared with NSCD there was an increase in IgG, and a decrease in IgA plasma cells, consistent with their potential role in CD fibrosis. Two Lumican-positive fibroblast subtypes were identified and subclassified based on expression of selectively enriched genes as fibroblast clusters (C)12 and C9. Cells within these clusters expressed the profibrotic genes *Decorin* (C12) and *JUN* (C9). C9 cells expressed *ACTA2*; ECM genes *COL4A1, COL4A2, COL15A1, COL6A3, COL18A1* and *ADAMDEC1*; *LAMB1* and *GREM1*. GO and KEGG Biological terms showed extracellular matrix, stricture organisation and regulation of *WNT* pathway genes are associated with the C12 and C9 gene sets. Trajectory and differential gene analysis of C12 and C9 identified four sub-clusters. Intra sub-cluster gene analysis detected co-regulated gene modules that aligned along predicted pseudotime trajectories and identified *CXCL14* and *ADAMDEC1* as key module markers. Our findings support further investigation of fibroblast heterogeneity and interactions with local and circulating immune cells at earlier time points in fibrosis progression. Breaking these interactions by targeting one or other population may improve therapeutic management for SCD.

## Introduction

Chronic inflammation is a significant factor in driving intestinal fibrosis and reciprocal interactions between fibroblasts and inflammatory immune cells are key to the pathogenesis of fibrosis.^1^ Stricture formation occurs in 30-50% of Crohn’s disease (CD) patients^2,3^ and stricturing CD (SCD) is linked with high levels of morbidity and healthcare utilisation.^4,5^ Fibrosis is a transmural process and current pharmacological treatments, including biologic and small molecule drugs, neither reduce fibrosis nor the requirement for stricture resection,^6^ and stricture recurrence is common.^7^

Fibrosis is characterised by increased deposition of extracellular matrix (ECM) proteins such as Collagen-I with accompanying thickening of both submucosa and smooth muscle cell layers.^8^ Fibroblasts are the primary source of ECM proteins, and both epithelial-to-mesenchymal transition (EMT) and the recruitment of circulating fibrocytes contribute to the intestinal fibroblast pool in SCD.^9-11^ The increase in ECM production in SCD is compounded by the reduced expression of enzymes that degrade collagen (matrix metalloproteinases (MMPs) and increased expression of tissue inhibitors of MMPs (TIMPs).^1^ The molecular mechanisms underlying SCD are complex, incompletely described and influenced by environmental triggers and genetic susceptibility^12-14^

Here, we report the use of human full thickness CD resection specimens and single cell RNA sequencing (scRNA-seq) to gain information on the cell types in the SCD and non-stricturing (NSCD) regions of the CD small bowel.

## Methods

### Single cell isolation from surgically resected tissue

Human full thickness CD resection specimens from the CD small bowel were washed with Hanks’ Balanced Salt solution (HBSS) supplemented with 0.01% Dithiothreitol and then HBSS-EDTA (1 mM) for 10 min per wash under agitation at 37°C. Finely dissected full thickness tissue was incubated with 20 mL of Dulbecco’s modified Eagle’s medium (DMEM) (Gibco) supplemented with collagenase type 1A (1 mg/mL, c2674-500 mg, Sigma) and DNase I (10 units/mL) for 45–60 min under gentle agitation, then filtered through a 100 μm cell strainer and the cell pellet washed twice in PBS. For scRNA-seq cells were re-suspended in 1 mL FBS supplemented with 10% DMSO) and cryostored in line with the recommended 10x genomic protocol for Fresh Frozen Human Peripheral Blood Mononuclear Cells.

### Single cell RNA sequencing

SCD and NSCD tissue was collected and processed independently (n=8 in total; SCD, 4 and NSCD 4 samples) as above.

Before library preparation, cell suspensions were thawed, washed (PBS BSA 0.04%), filtered and cell viability assessed; a viability cut-off of >70% was set for each sample to proceed. ScRNA-seq libraries were generated using the Chromium™ 3’ Library and Gel Bead Kit v3 (PN-1000092) (10X Genomics, California, USA). Final libraries were run on three NextSeq500 High-output v2.5 150-cycle kits (Illumina, CA) with a 26[8]98 cycle configuration to generate 50,000 reads per cell (10X Genomics recommendation).

Sequence reads were aligned with cellranger (v3.1.0; 10x genomics). Imported single cell gene counts were analysed using R package Seurat (v4). A data set was prepared per sample and each was normalised (function “NormalizeData”) followed by the identification of variable features (function “FindVariableFeatures” using parameters “vst”method and nfeatures =2000). The fraction of mitochondrial genes was calculated for quality assessment (function “PercentageFeatureSet”). Data sets were integrated by selecting integration features and anchors and then using the function “IntegrateData”. Integration data were then scaled (function “ScaleData”) and batch effects regressed out by incorporating patient ID. Principle component analysis (functions “RunPCA” and “FindNeighbors”) was used for graph based clustering at a resolution of 0.3 and Uniform Manifold Approximation (UMAP) and *t*-SNE dimensionality reduction was then computed (functions “RunUMAP”and “RunTSNE”). Batch correction was performed using Harmony by integrating the patient ID variable.^15^ Fibroblast and B clusters were identified by visualising cell markers (e.g., LUM and CD83 respectively) before being organised as independent sub-sets and analysed further. Monocle3 was used to subcluster and analyse pseudotime trajectories of B cell and fibroblast lineages.^16^ Other cell clusters were identified using scAnnotatR,^19^ as well as cross referencing markers provided in the single-cell type transcriptomics map and protein atlas of human tissues.^17-19^

Cross validation analysis of fibroblast cells (CD data set to the fibroblast atlas) used custom R scripts and marker gene identification (Seurat FindMarkers function) from both data sets. Tables of overlapping marker genes were assembled with the respective fold change metrics. Data imported into Cytoscape and StringDB networks were prepared using Omics visualizer app.^20^

### Ethics

Appropriate local Ethics Committee approvals (London - City Road & Hampstead Research Ethics Committee; 15/LO/2127) and informed consent were obtained prior to patient recruitment.

### Data Availability

The scRNA-seq data are available *via* Array Express accession code: E-MTAB-11792.

## Results

### Stricture-specific cell types in stricturing Crohn’s disease

We performed scRNA-seq of cells isolated from full-thickness tissue surgically resected from the strictured ileum of CD patients (n= 4) and non-strictured CD controls (NSCD n= 4); 3/4 were patient matched (Table 1). No prior selection was applied (e.g., FACS) to avoid cell type bias on sequencing. Instead, all cells were sequenced (total of 14,539 cells), integrated and batch corrected.^15^ UMAP visualised the complete integrated data set. This identified 24 clusters which were assigned to 16 definable cell lineages based on cell markers obtained from the published literature (Figure 1A and B and Supplementary Figure 1A).^21^ NSCD and SCD preparations contained the same cell types (Figure 2A and B) and the same proportions of endothelial and fibroblasts, indicating that phenotype did not impact cell release (Figure 2B; Supplementary Figure 1B and C).

**Table 1.**
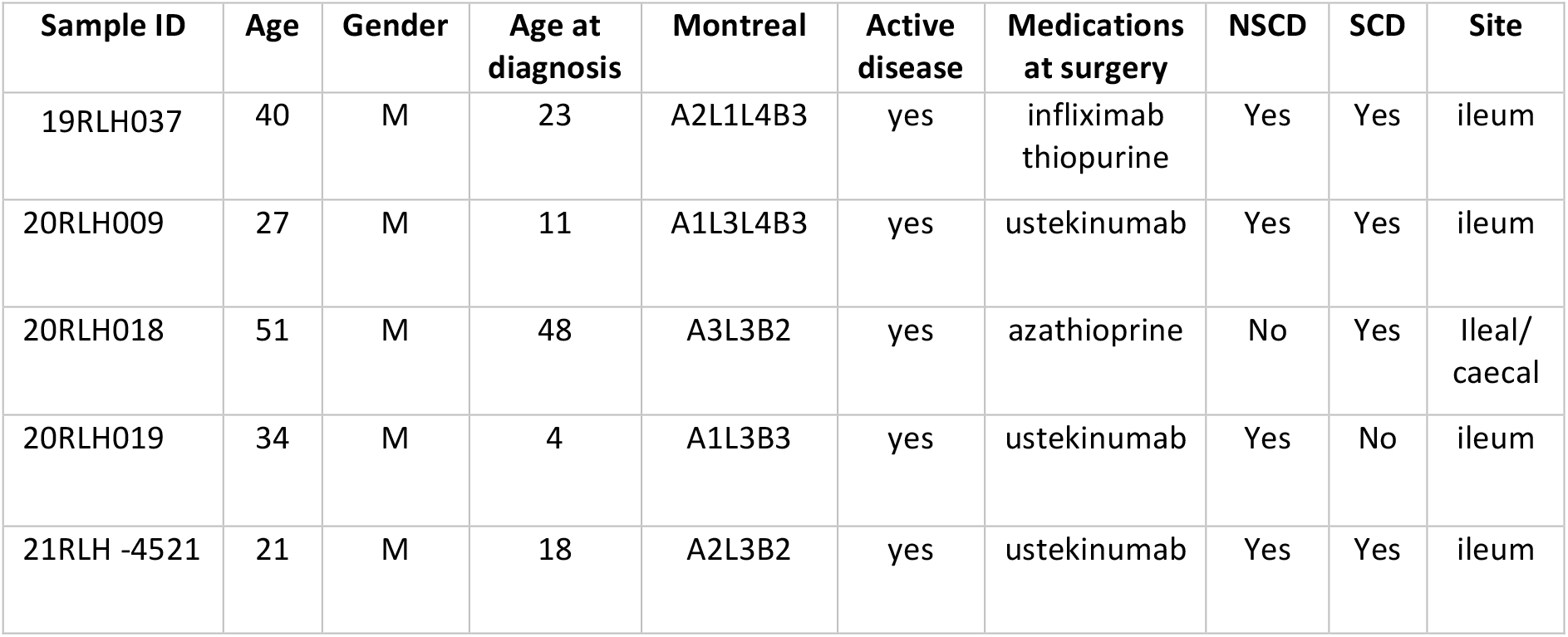
Clinical characteristics of Crohn’s disease cohort. All cases included in our study were “sporadic” Crohn’s disease (CD) with no cases of very early onset CD. This is evidenced by the fact that the disease duration at the time of surgery for the 2 paediatric diagnoses was 16 and 30 years. Several patients had a history of penetrating disease (Montreal B3), but resection specimens used in this analysis were harvested by a GI pathologist to ensure that they derived from fibrotic structures (SCD) or non strictured (NSCD) segments away from the site of any penetration.

| Sample ID | Age | Gender | Age at diagnosis | Montreal | Active disease | Medications at surgery | NSCD | SCD | Site |
| --- | --- | --- | --- | --- | --- | --- | --- | --- | --- |
| 19RLH037 | 40 | M | 23 | A2L1L4B3 | yes | infliximab<br>thiopurine | Yes | Yes | ileum |
| 20RLH009 | 27 | M | 11 | A1L3L4B3 | yes | ustekinumab | Yes | Yes | ileum |
| 20RLH018 | 51 | M | 48 | A3L3B2 | yes | azathioprine | No | Yes | Ileal/<br>caecal |
| 20RLH019 | 34 | M | 4 | A1L3B3 | yes | ustekinumab | Yes | No | ileum |
| 21RLH -4521 | 21 | M | 18 | A2L3B2 | yes | ustekinumab | Yes | Yes | ileum |

**Figure 1.**
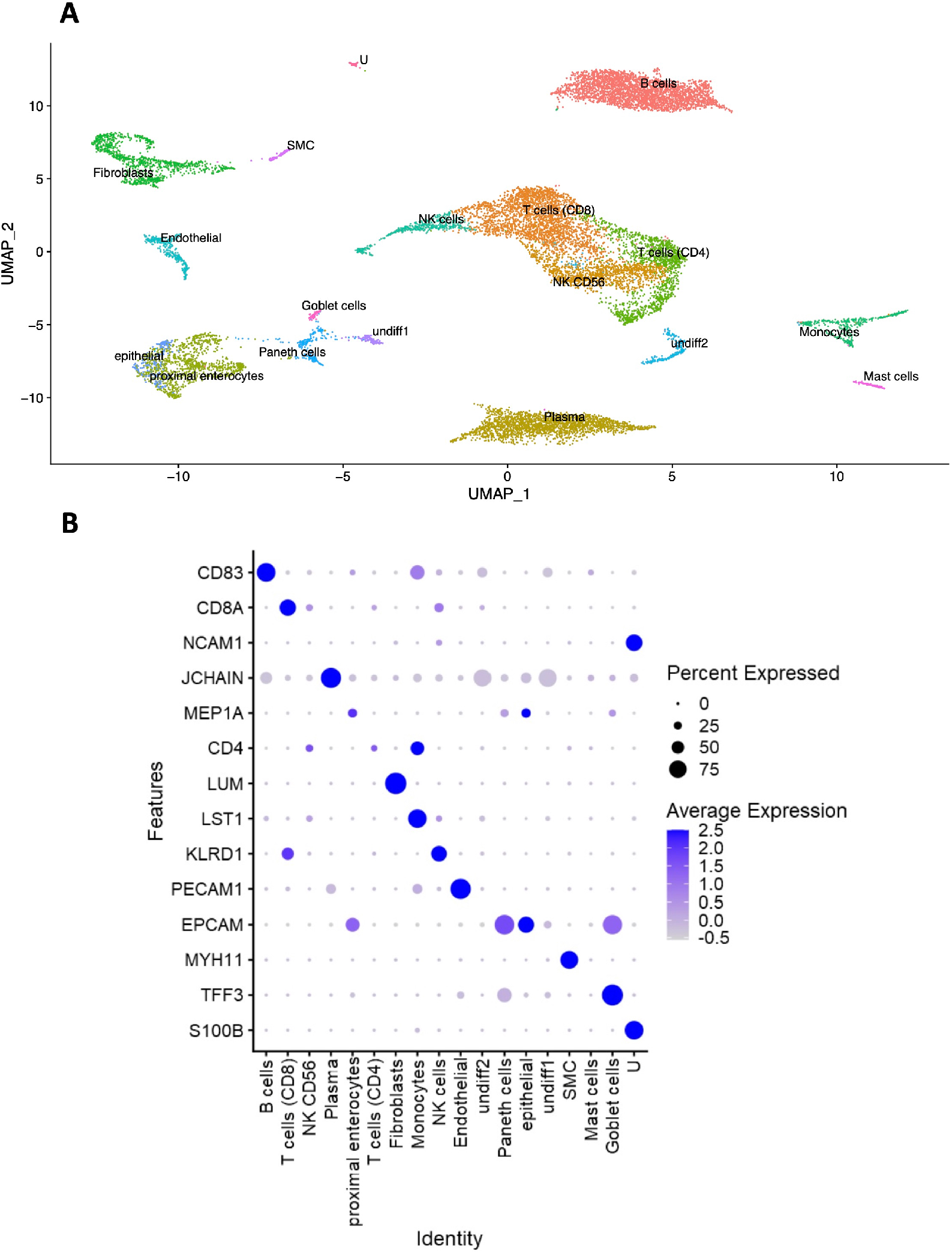
Cellular profile of stricturing Crohn’s disease (SCD). ScRNA-seq was performed on cells from full-thickness tissue surgically resected specimens from stricturing (SCD; n=4) and non-stricturing (NSCD; n=4) ileum of CD patients (n=5). Three patients provided both SCD and NSCD specimens. No prior cell selection was applied (e.g., FACs sorting) and all cells (n=14,539) were sequenced. **A**. Uniform Manifold Approximation (UMAP) identified 24 clusters assigned to 16 definable cell lineages (Supplementary Figure 1A). Undifferentiated cells (Undiff1/2) contained markers of undifferentiated cells as defined by protein atlas webtool,^20^ and expressed TOP2A and MKI67 unlike any other cell type. Unknown (U) had no identifiable gene marker linked to any cell type. **B**. Bubble plot shows levels of expression of key markers for different cell types identified from published data sets.

**Figure 2.**
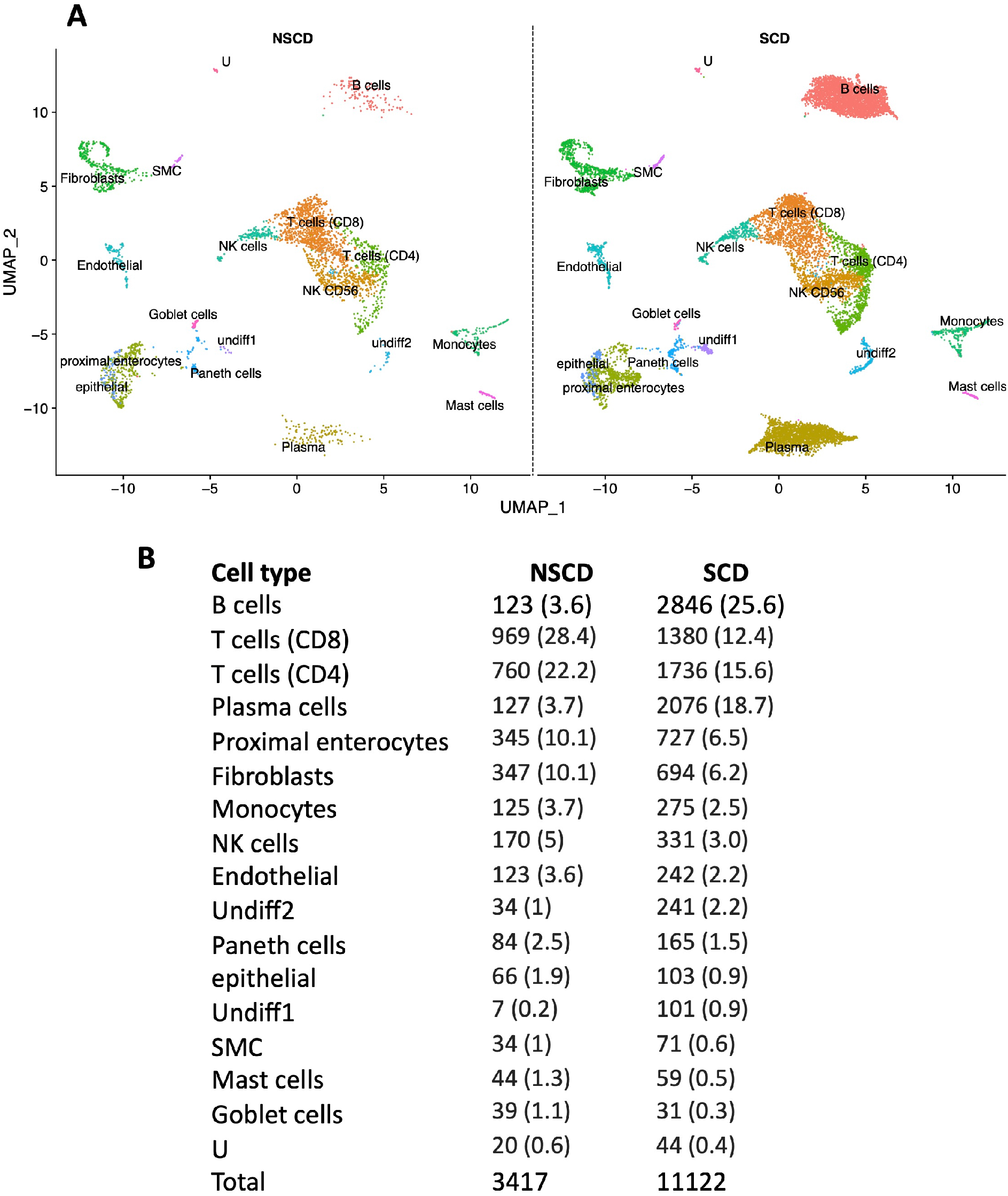
A. Stricturing (SCD) and non-stricturing (NSCD) small bowel tissue comprise the same cell types but proportions differ markedly for B cells and plasma cells. **A** UMAP of NSCD and SCD preparations indicate they are comprised of the same cell types. **B**. Numbers and frequency (%) of cells identified from NSCD and SCD small bowel tissue. B cells and plasma cells showed the most marked increased in frequency in SCD compared to NSCD.

### Stricturing Crohn’s disease is associated with selective increases in B cells and IgG plasma cells

Fibroblast function under inflammatory conditions is shaped by interaction with immune cells.^1^ Amongst immune cell types, B cells and plasma cells were most markedly increased in frequency in SCD compared to NSCD (Figure 2A and B). As well as differences in absolute numbers of plasma cells, there were differences in the distribution of antibody subclasses produced by these cells. The proportion of IgG subclasses was increased in SCD, whereas IgA2, was increased in NSCD (Figure 3A). *CXCR4*, which is reported to facilitate plasma cell homing to inflamed tissue was expressed by 62% of plasma cells in SCD *versus* 44% in NSCD (Figure 3B).^22^ Additional sub-clustering of B cells identified a population of naïve B cells (Figure 3C, cluster 1), expressing *IGHD* and *IGHM*; a cluster expressing genes consistent with a memory B cell phenotype (Figure 3C, cluster 2); and a small subset of B cells expressing germinal centre markers including *BCL6* and *AICDA* (Figure 3C, cluster 3), the latter of which were almost exclusively found in SCD but not NSCD.

**Figure 3.**
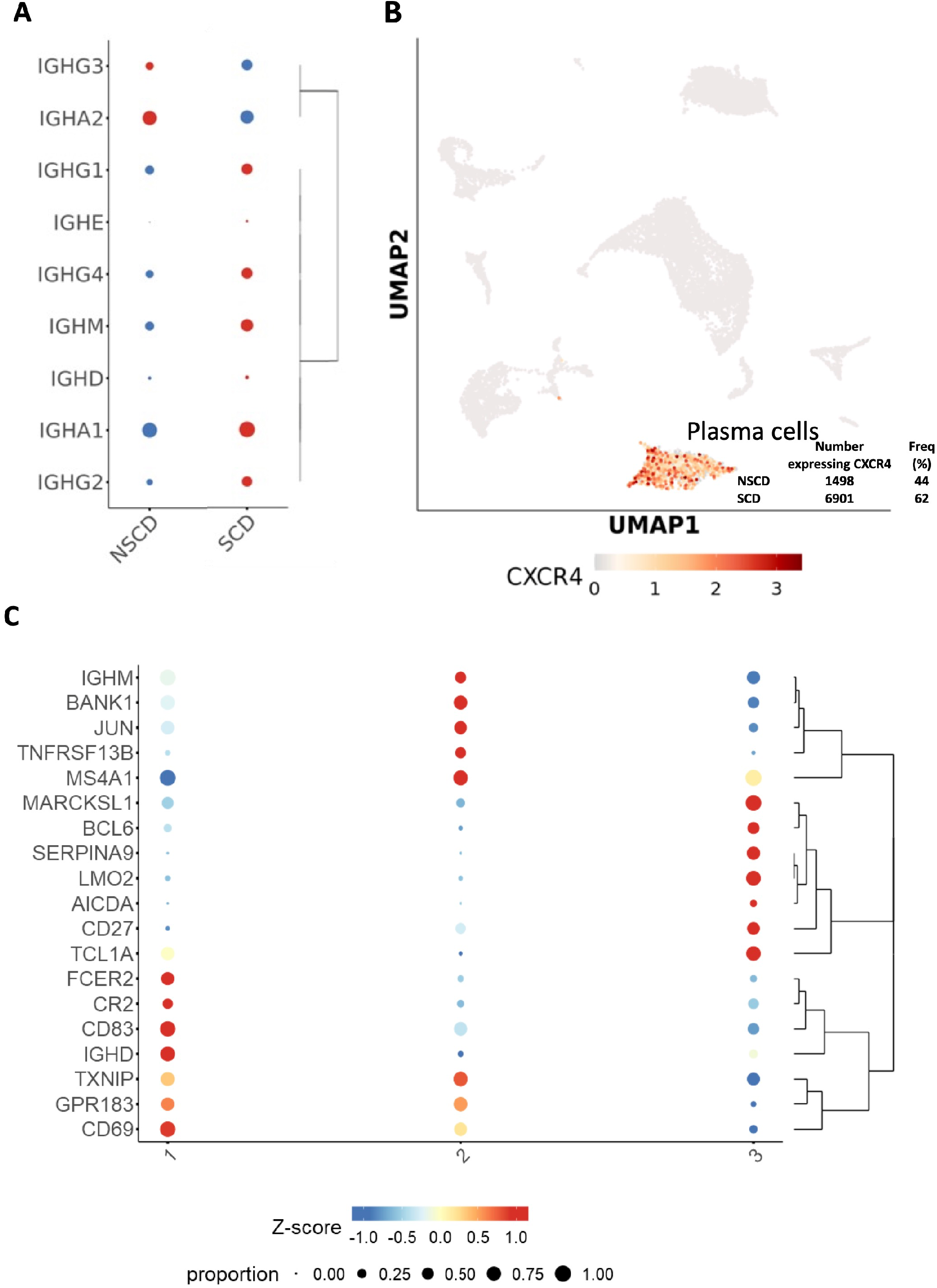
Analysis of B cells and plasma cells in SCD and NSCD small bowel tissue. **A**. Proportions of antibody subclasses expressed by plasma cells in SCD and NSCD tissues. **B**. Expression of *CXCR4* in plasma cells. **C**. Sub-clustering of B cells identified three distinct clusters consistent with naïve (1), memory (2) and germinal centre (3) subsets.

### Identification of two fibroblast subtypes

Two Lumican (LUM)+ve fibroblast subtypes were identified from our scRNA-seq data (Figure 1A and B). These were further subclassified as two clusters (fibroblast cluster 12 (C12) and fibroblast cluster 9 (C9)) based on expression of selectively enriched genes (fold change >2 and adjusted P-value ≤0.001; (Supplementary Figure 1D and E; Supplementary Table 1). Ranking these genes according to the numbers of cells in which they were expressed demonstrated that the highest proportion of cells within the two clusters expressed the profibrotic genes *Decorin* (*DCN*) for C12 and *JUN* for C9. Cells in C9 also expressed *ACTA2*; the ECM genes *COL4A1, COL4A2, COL15A1, COL6A3, COL18A1* and *ADAMDEC1*; as well as *LAMB1* and *GREM1* (Supplementary Table 1).

### Extracellular matrix, stricture organisation and regulation of WNT pathway genes are associated with fibroblast clusters

The GO and KEGG Biological terms associated with the fibroblast cluster C12 and C9 gene sets included ECM and stricture organisation (both clusters; GO terms), and positive and negative regulation of *WNT* pathway genes (C9; GO terms) (Supplementary Figure 2A-H). The fibroblast subtypes corresponded to those identified previously by scRNA-seq in ulcerative colitis (UC) and other fibrotic diseases.^23,24^ C9 fibroblast markers (Supplementary Table 2) were validated independently by comparing with PI16 fibroblasts in the perturbed human fibroblast atlas (fibroXplorer database;^24^ n=3,604 fibroblasts; (Supplementary Figure 3A and B). Differentiation trajectories of the two fibroblast populations were analysed,^25^ and four sub-clusters resolved (Supplementary Figure 3C). Differential gene analysis identified gene markers for each sub-cluster and intra sub-cluster gene analysis identified co-regulated gene modules that aligned along predicted pseudotime trajectories (Supplementary Figure 3D and E). The top two markers of module1 were *CXCL14* and *ADAMDEC1* (Supplementary Figure 4), previously identified to be markers in a distinct fibroblast population in UC and other fibrotic diseases,^23^ indicating that sub-cluster 1 are the same *ADAMDEC1*-like fibroblasts.

## Discussion

We have, for the first time, generated sequencing data from all the cell types isolated from resected full-thickness SCD and NSCD intestinal specimens from stricturing CD patients. Access to well phenotyped, fresh surgical specimens, rather than the more readily accessible mucosal biopsies, allowed sampling from the deeper layers of the bowel that contribute to CD fibrosis. This underlines the clinical importance of our dataset. In contrast to studies in other organ systems that have used enrichment, we determined natural groupings from an atlas of 14,539 sequenced cells. This identified fibroblasts and immune cell populations in the gut without any potential bias associated with pre-selection. The availability of human IBD datasets, such as fibroXplorer,^24^ supported independent validation.

The numbers of B cells and plasma cells were increased in SCD tissue to a much greater degree than other immune cell populations. Several studies have recently identified a role for B cells in the pathogenesis of UC,^22,26,27^ but their contribution to CD remains poorly understood. There was a shift towards plasma cells expressing IgG subclasses, and away from IgA, in SCD. IgG drives inflammation in UC by activating innate immune cells *via* interactions with Fc receptors and the consequent promotion of inflammatory Th17 responses.^27^ In UC, IgG plasma cells infiltrate the inflamed intestine *via* CXCR4,^22^ and this chemokine receptor was expressed by many plasma cells in our CD study. IgG plasma cells expressing *CXCR4* were enriched in SCD compared with NSCD samples. Reclustering of B cells revealed naïve, memory and germinal centre B cell populations, the latter of which were almost exclusively found in SCD. These finding are consistent with the formation of organised tertiary lymphoid tissue (TLT) in SCD.^28^ TLT formation is a feature of CD,^29^ but its causal contribution to the inflammatory process or to fibrosis is currently poorly understood. In other inflammatory contexts, B cells have been shown to interact directly with fibroblasts to influence the fibrotic process in inflammation,^30^ raising the possibility that a similar process contributes to fibrosis in CD.

We identified two LUM+ve fibroblast subtypes, which were further subclassified as two clusters, C12 and C9. Their analysis highlighted the expression of profibrotic genes *DCN* and *JUN* (C12 and C9, respectively): DCN modulates TGFβ-driven fibrosis in a number of organs and JUN induction drives murine pulmonary fibrosis.^31-33^ Expression of *ACTA2* (C9), as well as ECM genes including *COL4A2, ADAMDEC1, LAMB1* and *GREM1* was found. Interestingly, *LAMB1* and *COL4A2* are potential markers of fibrosis progression in liver and *GREM1*, a highly conserved member of the TGFβ superfamily, is involved in fibrosis across multiple organ systems.^34,35^

Our study provides an unbiased reference dataset for SCD *versus* NCD, which is controlled internally because most of these tissues are matched from the same individual. This enables the internal relationships of different cell types to be compared in a single time window. Our data indicate clearly that therapeutic intervention must come much earlier in CD progression than the timepoint described here to break the apparent interaction between immune cells and fibroblasts. In this context, it would be interesting to consider whether increased immune cells residency in the mucosa and/ or assay of circulating antibodies/ immune cells in blood could indicate fibrotic progression at a much earlier time point.

## Abbreviations

CD: Crohn’s disease
DMEM: Dulbecco’s modified Eagle medium
ECM: extracellular matrix
C12: fibroblast cluster 12
C9: fibroblast cluster 9
H3K27ac: histone-3 acetylation at lysine 27
HDAC: histone deacetylase
OHPro: hydroxyproline
IF: Immunofluorescence
IBD: inflammatory bowel disease
LME: linear mixed effects model
MSigDB: Molecular Signatures Database
NSCD: non-stricturing CD
PCs: principal components
PCA: Principal components analysis
QC: quality control
scATAC-seq: scAssay for Transposase-Accessible Chromatin using sequencing
scRNA-seq: single cell RNA sequencing
SCD: stricturing CD
UC: ulcerative colitis
UMAP: Uniform Manifold Approximation
VPA: valproic acid.

## Acknowledgements

The authors thank participating patients and members of the Blizard Institutes’ core facilities.

## Supplementary Figures

**Supplementary Figure 1.**
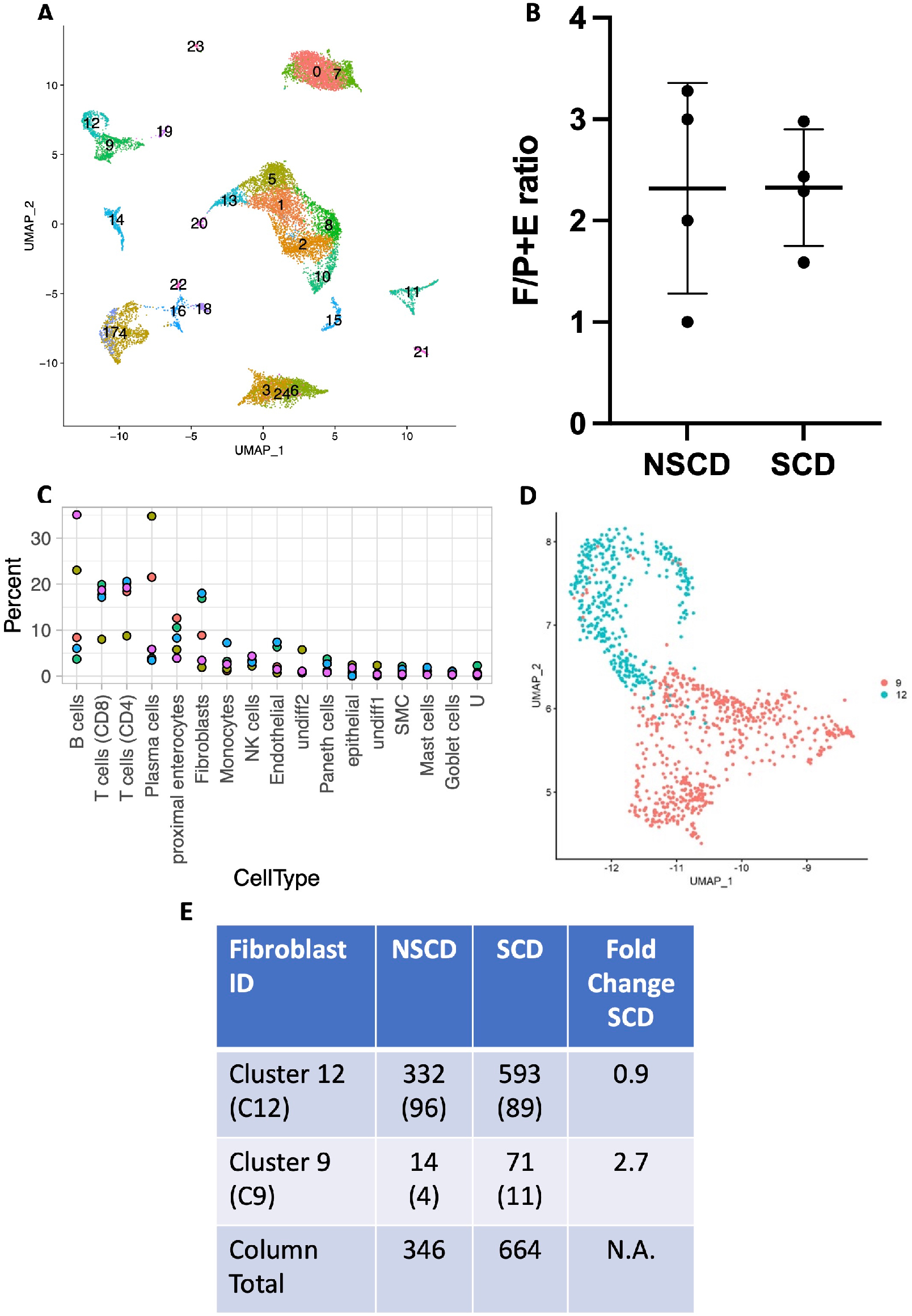
Schematic of single cell sequencing process and identification of fibroblast subtypes from SCD and NSCD small bowel tissue. **A**. UMAP showing marking each of the clusters identified using the Seurat pipeline (see also Figure 1A). **B**. Comparison of fibroblast (F) cell numbers with pericytes plus endothelial (P+E) cell numbers from resected SCD and NSCD tissue showed no significant differences suggesting consistent release of cells across both stricturing and non-stricturing tissue. **C**. Comparison of portion of cell types identified for each processed scRNA-seq specimen where each dot represents a particular resected specimen. The proportion of epithelial cells was as expected and explained by the washing efficiency during tissue collection. **D**. UMAP of fibroblasts identifying C12 and C9 fibroblast clusters. **E**. Comparison of the numbers of LUM+ve fibroblasts in cluster 12 (C12) and cluster 9 (C9) in resected small bowel NSCD and SCD tissue. The percentage of each cluster is given in parenthesis and the fold change in the proportion of C12 and C9 fibroblasts in NSCD and SCD is shown relative to SCD.

**Supplementary Figure 2.**
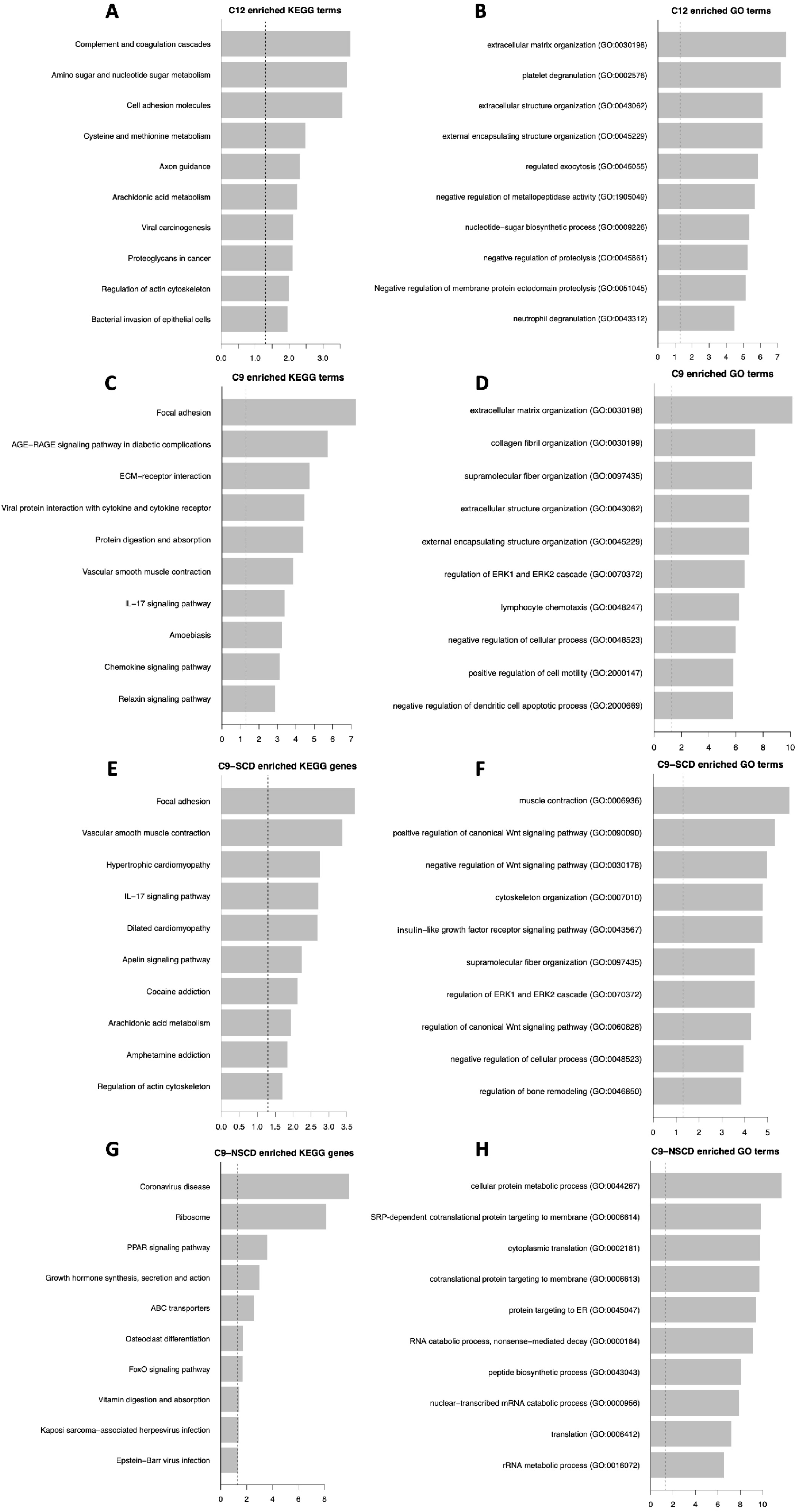
Enriched Kegg and GO terms for fibroblast clusters C12 and C9. Top 10 enriched KEGG and GO terms for C12 fibroblasts (**A** and **B** respectively), C9 fibroblasts (**C** and **D** respectively). Gene signatures from C9 fibroblasts were further analysed by identifying the top 10 enriched KEGG and GO terms of C9 fibroblasts originating from SCD (**E** and **F** respectively) or NSCD (**G** and **H** respectively) conditions.

**Supplementary Figure 3.**
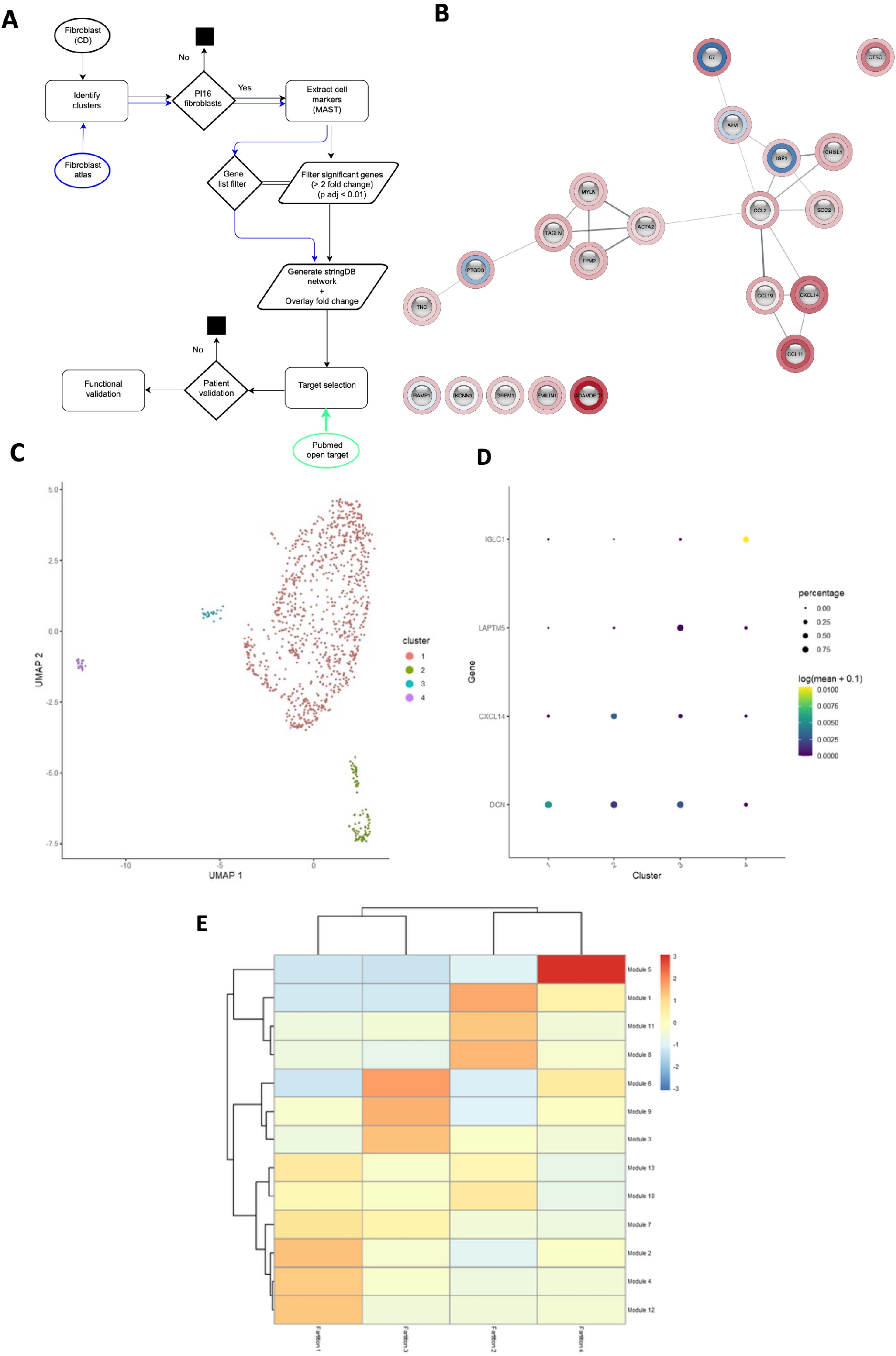
Workflow and stringdb network assembly, trajectory and differential gene analysis for fibroblast populations and clusters. **A**. Schematic of scRNA-seq workflow, incorporating stringdb network assembly for C9-like fibroblasts and target validation workflow. **B**. Stringdb network for C9-like SCD fibroblasts. **C**. Four clusters resolved from differentiation trajectories of the two fibroblast populations were analysed using Monocle3.^16^ **D**. Identification of the most significant gene marker for each cluster using differential gene analysis. **E**. Co-regulated gene modules identified from predicted pseudotime trajectories using Monocle3.^16^ Each module contains multiple genes that have similar expression patterns between cell neighbours (i.e., intracluster gene analysis). Gene members of each gene module are listed in Supplementary Figure 4.

**Supplementary Figure 4.**
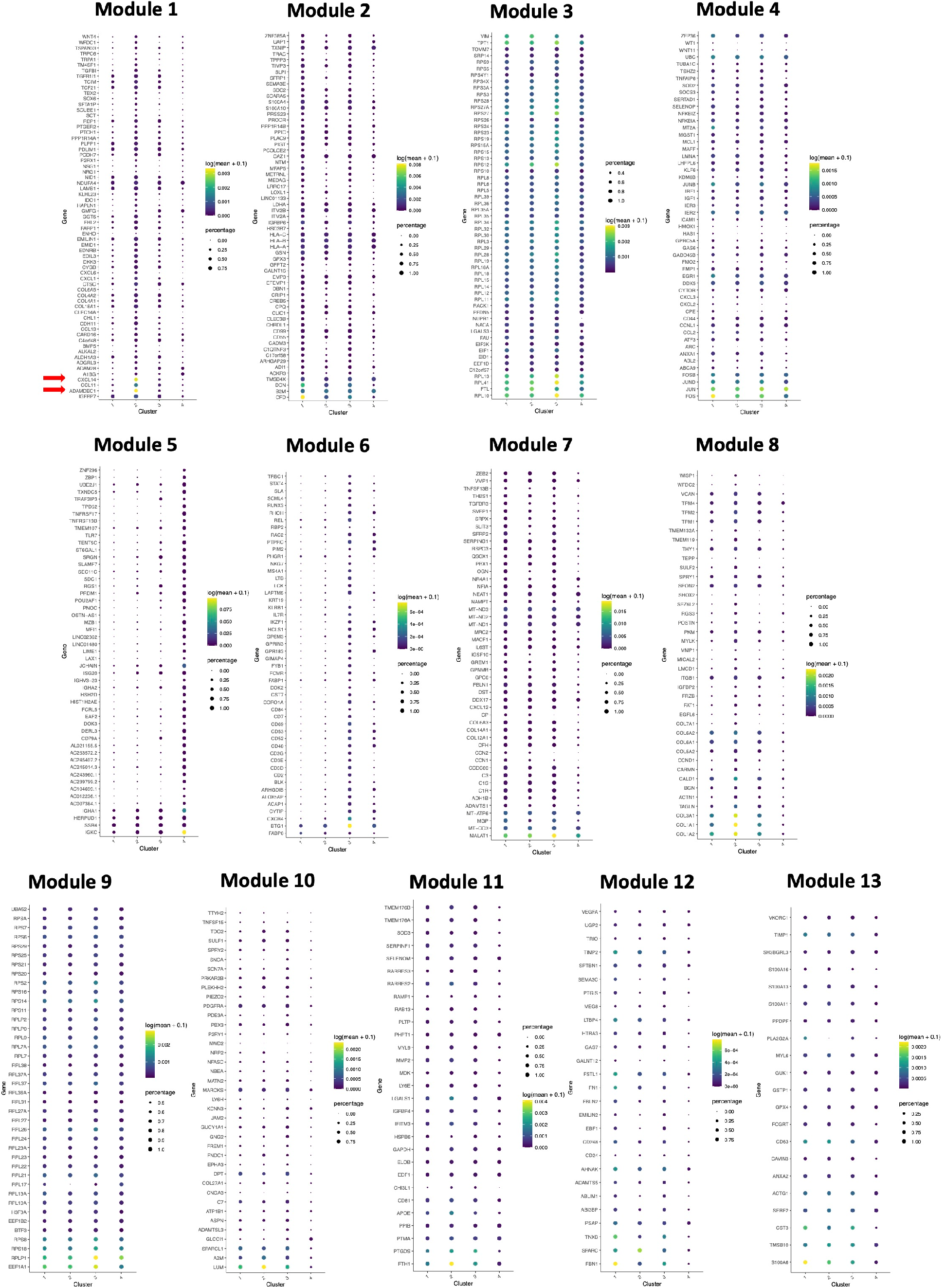
Gene markers for co-regulated modules. All gene modules were identified using Monocle3,^16^ as described in Figure 3. Note that *CXCL14* and *ADAMDEC1* were the top two markers of module1 (arrowed), and are primarily expressed in sub-cluster2

