## Supplementary tables 1 and 2 for "Single cell sequencing data identify distinct B cell and fibroblast populations in stricturing Crohn’s disease"

**Supplementary Table 1. Gene markers that define fibroblast cell clusters C9 and C12.** List of all differentially expressed genes identified using the findmarkers function (MAST algorithm) to compare C9 to C12 fibroblast populations. Each table has been ordered on pct.1 and pct.2 columns.

**C12 compared to C9**

| Gene | p_val | avg_log2FC | pct.1 | pct.2 | p_val_adj |
|---|---|---|---|---|---|
| DCN | 5.52E-39 | 0.727796 | 1 | 0.949 | 1.4E-34 |
| CFD | 1.2E-54 | 1.112183 | 0.997 | 0.858 | 3.03E-50 |
| TMSB4X | 9.7E-36 | 0.781292 | 0.997 | 0.961 | 2.45E-31 |
| CST3 | 2.02E-47 | 0.734969 | 0.997 | 0.922 | 5.11E-43 |
| B2M | 6.59E-15 | 0.627643 | 0.997 | 0.986 | 1.67E-10 |
| IGFBP6 | 9.5E-128 | 2.058368 | 0.995 | 0.748 | 2.4E-123 |
| PLAC9 | 3E-123 | 1.766458 | 0.995 | 0.638 | 7.6E-119 |
| S100A10 | 2.2E-110 | 1.583371 | 0.995 | 0.757 | 5.7E-106 |
| HLA-B | 1.07E-19 | 0.648267 | 0.995 | 0.942 | 2.71E-15 |
| FBN1 | 2.4E-120 | 2.136493 | 0.992 | 0.792 | 6E-116 |
| S100A6 | 3.48E-66 | 1.047138 | 0.992 | 0.938 | 8.79E-62 |
| GSN | 2.29E-38 | 0.769769 | 0.992 | 0.883 | 5.8E-34 |
| TIMP1 | 2.42E-57 | 1.078245 | 0.99 | 0.852 | 6.11E-53 |
| TIMP2 | 9.52E-31 | 0.611803 | 0.99 | 0.863 | 2.41E-26 |
| TNXB | 7.22E-82 | 1.421838 | 0.987 | 0.76 | 1.82E-77 |
| S100A4 | 3.38E-83 | 1.312589 | 0.987 | 0.815 | 8.56E-79 |
| TIMP3 | 1.81E-63 | 1.175705 | 0.975 | 0.711 | 4.59E-59 |
| FSTL1 | 5.46E-71 | 1.470796 | 0.97 | 0.747 | 1.38E-66 |
| ANXA2 | 2.01E-55 | 0.951132 | 0.97 | 0.776 | 5.08E-51 |
| LTBP4 | 3.56E-47 | 0.885592 | 0.97 | 0.77 | 9E-43 |
| MFAP5 | 8.3E-162 | 2.944869 | 0.957 | 0.297 | 2.1E-157 |
| SPARC | 2.14E-18 | 0.620161 | 0.957 | 0.81 | 5.42E-14 |
| EFEMP1 | 5.63E-49 | 0.952892 | 0.952 | 0.698 | 1.42E-44 |
| CD248 | 9.05E-95 | 1.524008 | 0.95 | 0.477 | 2.29E-90 |
| S100A11 | 1.69E-40 | 0.893673 | 0.947 | 0.742 | 4.26E-36 |
| IGFBP5 | 1.04E-16 | 0.734037 | 0.942 | 0.779 | 2.64E-12 |
| FN1 | 4.11E-93 | 2.09181 | 0.92 | 0.417 | 1.04E-88 |
| EMP3 | 1.06E-56 | 1.192682 | 0.92 | 0.653 | 2.67E-52 |
| CD99 | 5.59E-30 | 0.653976 | 0.915 | 0.751 | 1.41E-25 |
| TXNIP | 5.17E-18 | 0.619232 | 0.915 | 0.748 | 1.31E-13 |
| FBLN2 | 3.51E-84 | 1.493094 | 0.912 | 0.386 | 8.88E-80 |
| GPX3 | 2.46E-32 | 0.748008 | 0.91 | 0.631 | 6.21E-28 |
| TAGLN2 | 1.48E-25 | 0.590009 | 0.905 | 0.661 | 3.74E-21 |
| SEMA3C | 3E-139 | 2.39509 | 0.894 | 0.179 | 7.5E-135 |
| CD55 | 8.5E-116 | 2.439183 | 0.889 | 0.333 | 2.1E-111 |
| PCOLCE | 3.05E-19 | 0.614978 | 0.879 | 0.672 | 7.72E-15 |
| SFRP1 | 1.5E-23 | 0.698172 | 0.877 | 0.602 | 3.8E-19 |
| RHOA | 1.23E-27 | 0.69305 | 0.869 | 0.652 | 3.11E-23 |

| | | | | | |
|---|---|---|---|---|---|
| RNH1 | 1.11E-25 | 0.620559 | 0.867 | 0.625 | 2.81E-21 |
| CPQ | 5.83E-30 | 0.638886 | 0.862 | 0.572 | 1.47E-25 |
| SDC2 | 7.94E-44 | 0.883742 | 0.859 | 0.465 | 2.01E-39 |
| CLU | 6.32E-34 | 0.819867 | 0.854 | 0.502 | 1.6E-29 |
| SCARA5 | 1.9E-106 | 1.820405 | 0.849 | 0.205 | 4.9E-102 |
| SH3BGRL3 | 8.67E-56 | 1.267391 | 0.847 | 0.513 | 2.19E-51 |
| LDHA | 2.64E-21 | 0.666494 | 0.839 | 0.614 | 6.67E-17 |
| C1QTNF3 | 3.25E-34 | 0.726573 | 0.827 | 0.451 | 8.21E-30 |
| MT2A | 1.92E-11 | 1.002204 | 0.824 | 0.636 | 4.85E-07 |
| PLA2G2A | 1.96E-56 | 1.757877 | 0.822 | 0.348 | 4.95E-52 |
| CLIC1 | 9.11E-21 | 0.610611 | 0.819 | 0.593 | 2.3E-16 |
| UAP1 | 5.87E-73 | 1.5602 | 0.809 | 0.305 | 1.48E-68 |
| VKORC1 | 2.64E-21 | 0.598223 | 0.804 | 0.591 | 6.67E-17 |
| PPIC | 1.21E-47 | 1.074767 | 0.796 | 0.437 | 3.06E-43 |
| SMIM14 | 1.67E-26 | 0.63365 | 0.779 | 0.456 | 4.21E-22 |
| ADAMTS5 | 2.71E-80 | 1.992449 | 0.774 | 0.23 | 6.84E-76 |
| PIGT | 4.54E-39 | 0.843917 | 0.771 | 0.393 | 1.15E-34 |
| CADM3 | 2.98E-72 | 1.416784 | 0.769 | 0.232 | 7.53E-68 |
| HTRA3 | 1.25E-34 | 1.215896 | 0.766 | 0.415 | 3.17E-30 |
| C17orf58 | 8.71E-81 | 1.80359 | 0.764 | 0.226 | 2.2E-76 |
| CLEC3B | 1E-114 | 1.844727 | 0.759 | 0.1 | 2.6E-110 |
| KLF2 | 2.29E-13 | 0.732776 | 0.746 | 0.572 | 5.78E-09 |
| CLTB | 2.38E-30 | 0.795704 | 0.744 | 0.417 | 6.01E-26 |
| MEDAG | 1.4E-34 | 0.795629 | 0.741 | 0.353 | 3.55E-30 |
| ABLIM1 | 2.51E-42 | 0.871546 | 0.739 | 0.309 | 6.35E-38 |
| CD34 | 1.3E-38 | 0.858977 | 0.739 | 0.336 | 3.29E-34 |
| TPPP3 | 2.77E-67 | 1.562293 | 0.734 | 0.232 | 7.01E-63 |
| GAS7 | 3.09E-44 | 1.041554 | 0.734 | 0.331 | 7.82E-40 |
| REXO2 | 6.31E-18 | 0.597582 | 0.731 | 0.496 | 1.6E-13 |
| ACKR3 | 3.69E-73 | 1.557218 | 0.724 | 0.179 | 9.34E-69 |
| CREB5 | 3E-47 | 1.099362 | 0.721 | 0.281 | 7.59E-43 |
| TRIOBP | 5.68E-30 | 0.670557 | 0.721 | 0.365 | 1.44E-25 |
| METRNL | 1.11E-63 | 1.259878 | 0.714 | 0.227 | 2.81E-59 |
| PTGIS | 1.95E-37 | 0.903776 | 0.711 | 0.313 | 4.93E-33 |
| DBN1 | 8.1E-69 | 1.243779 | 0.686 | 0.17 | 2.05E-64 |
| SERPINE2 | 1.47E-19 | 0.887899 | 0.686 | 0.4 | 3.72E-15 |
| ADI1 | 5.91E-26 | 0.739161 | 0.686 | 0.376 | 1.5E-21 |
| ITM2A | 3.27E-41 | 1.135903 | 0.678 | 0.291 | 8.26E-37 |
| ZNF385A | 3.06E-49 | 1.031576 | 0.673 | 0.229 | 7.73E-45 |
| ARHGAP29 | 2.77E-46 | 0.941716 | 0.666 | 0.23 | 7.01E-42 |
| UGP2 | 2.22E-22 | 0.763663 | 0.653 | 0.381 | 5.62E-18 |
| OSR2 | 3.58E-29 | 0.757793 | 0.651 | 0.292 | 9.04E-25 |
| CRIP1 | 6.17E-29 | 1.217787 | 0.641 | 0.302 | 1.56E-24 |
| YWHAH | 4.97E-25 | 0.772146 | 0.631 | 0.316 | 1.26E-20 |
| PRSS23 | 5.89E-59 | 1.259388 | 0.628 | 0.156 | 1.49E-54 |
| AXL | 5.19E-25 | 0.67752 | 0.621 | 0.305 | 1.31E-20 |

| | | | | | |
|---|---|---|---|---|---|
| F10 | 1.33E-19 | 0.666597 | 0.621 | 0.336 | 3.36E-15 |
| PHGDH | 1.03E-38 | 0.755862 | 0.613 | 0.21 | 2.6E-34 |
| SLPI | 5.68E-52 | 1.76331 | 0.595 | 0.16 | 1.44E-47 |
| PI16 | 3.24E-68 | 1.864965 | 0.59 | 0.09 | 8.2E-64 |
| PCOLCE2 | 4.2E-79 | 1.823331 | 0.59 | 0.072 | 1.06E-74 |
| HEG1 | 1.03E-29 | 0.589509 | 0.583 | 0.23 | 2.6E-25 |
| PPP1R14B | 2.09E-23 | 0.751395 | 0.575 | 0.292 | 5.28E-19 |
| LOXL1 | 7.85E-38 | 1.005406 | 0.573 | 0.199 | 1.99E-33 |
| CLDN11 | 6.2E-29 | 0.634452 | 0.573 | 0.226 | 1.57E-24 |
| LRRC17 | 5.3E-49 | 1.05657 | 0.568 | 0.134 | 1.34E-44 |
| SEMA3E | 1.92E-67 | 1.371876 | 0.555 | 0.072 | 4.86E-63 |
| UGDH | 3.35E-32 | 0.72396 | 0.553 | 0.194 | 8.47E-28 |
| LINC01133 | 2.04E-52 | 1.01731 | 0.535 | 0.104 | 5.17E-48 |
| ADAMTSL4 | 7.8E-38 | 0.693549 | 0.535 | 0.152 | 1.97E-33 |
| EMILIN2 | 6.67E-47 | 0.846781 | 0.528 | 0.114 | 1.69E-42 |
| GFPT2 | 4.76E-29 | 0.774514 | 0.513 | 0.179 | 1.2E-24 |
| VASN | 1.6E-27 | 0.714992 | 0.508 | 0.182 | 4.04E-23 |
| PROCR | 1.21E-45 | 1.105508 | 0.492 | 0.1 | 3.07E-41 |
| TBC1D12 | 5.34E-27 | 0.652958 | 0.492 | 0.173 | 1.35E-22 |
| ITGA11 | 5.87E-34 | 0.789623 | 0.48 | 0.131 | 1.48E-29 |
| SEMA3B | 1.33E-37 | 0.743557 | 0.477 | 0.115 | 3.37E-33 |
| CYTOR | 3.67E-19 | 0.724797 | 0.467 | 0.201 | 9.28E-15 |
| TRIO | 1.12E-21 | 0.727835 | 0.455 | 0.179 | 2.84E-17 |
| HSD3B7 | 4.39E-39 | 0.683122 | 0.437 | 0.086 | 1.11E-34 |
| C12orf75 | 1.26E-36 | 0.711187 | 0.435 | 0.095 | 3.18E-32 |
| SHISA3 | 4.96E-19 | 0.638221 | 0.417 | 0.159 | 1.25E-14 |
| GALNT15 | 2.31E-39 | 0.891362 | 0.384 | 0.058 | 5.85E-35 |
| AIF1L | 5.59E-46 | 0.696697 | 0.384 | 0.04 | 1.41E-41 |
| TNFAIP6 | 3.15E-07 | 0.69611 | 0.379 | 0.23 | 0.007957 |
| GALNT12 | 1.29E-32 | 0.653758 | 0.372 | 0.07 | 3.25E-28 |
| NTM | 1.23E-27 | 0.597367 | 0.334 | 0.067 | 3.11E-23 |
| DPP4 | 3.63E-25 | 0.593136 | 0.329 | 0.076 | 9.18E-21 |
| TRAC | 3.14E-30 | 0.670273 | 0.279 | 0.034 | 7.95E-26 |
| ADAMTS16 | 8.99E-30 | 0.636292 | 0.259 | 0.026 | 2.27E-25 |
| CD24 | 3.4E-19 | 0.772708 | 0.239 | 0.048 | 8.6E-15 |
| PRG4 | 2.2E-09 | 0.642053 | 0.048 | 0 | 5.57E-05 |

**C9 compared to C12**

| | p_val | avg_log2FC | pct.1 | pct.2 | p_val_adj |
|---|---|---|---|---|---|
| JUN | 2.35E-23 | 0.891514 | 0.947 | 0.962 | 5.94E-19 |
| EGR1 | 2.56E-09 | 0.660965 | 0.908 | 0.894 | 6.47E-05 |
| CALD1 | 1.09E-33 | 0.65643 | 0.9 | 0.965 | 2.75E-29 |
| SPARCL1 | 3.85E-35 | 0.883563 | 0.897 | 0.905 | 9.73E-31 |
| FOSB | 1.13E-26 | 0.961446 | 0.885 | 0.749 | 2.87E-22 |
| CXCL12 | 1.17E-32 | 0.943173 | 0.876 | 0.789 | 2.95E-28 |

| | | | | | |
|---|---|---|---|---|---|
| COL6A3 | 7E-18 | 0.652227 | 0.857 | 0.872 | 1.77E-13 |
| DPT | 1.48E-16 | 0.847522 | 0.788 | 0.714 | 3.74E-12 |
| SAT1 | 1.09E-09 | 0.753806 | 0.737 | 0.739 | 2.75E-05 |
| SOD2 | 4.78E-18 | 1.170726 | 0.72 | 0.643 | 1.21E-13 |
| PPP1R15A | 4.15E-13 | 0.659816 | 0.709 | 0.651 | 1.05E-08 |
| PTGDS | 1.56E-26 | 1.483627 | 0.687 | 0.394 | 3.94E-22 |
| APOE | 1.8E-09 | 0.727634 | 0.687 | 0.565 | 4.55E-05 |
| BTG1 | 1.67E-18 | 0.846339 | 0.683 | 0.595 | 4.22E-14 |
| IGFBP7 | 5.6E-12 | 0.680825 | 0.683 | 0.764 | 1.42E-07 |
| LHFPL6 | 3.54E-17 | 0.599089 | 0.67 | 0.686 | 8.96E-13 |
| C7 | 1.33E-51 | 2.38598 | 0.669 | 0.239 | 3.36E-47 |
| A2M | 4.76E-27 | 1.169691 | 0.663 | 0.568 | 1.2E-22 |
| IGF1 | 2.81E-21 | 1.023047 | 0.644 | 0.497 | 7.1E-17 |
| TPM2 | 1.36E-40 | 1.432498 | 0.641 | 0.41 | 3.44E-36 |
| RSPO3 | 5.45E-22 | 0.635414 | 0.636 | 0.671 | 1.38E-17 |
| LAMB1 | 3.91E-13 | 0.604513 | 0.599 | 0.548 | 9.9E-09 |
| GREM1 | 8.39E-15 | 1.107837 | 0.565 | 0.344 | 2.12E-10 |
| CRISPLD2 | 3.11E-14 | 0.694896 | 0.561 | 0.47 | 7.87E-10 |
| EMILIN1 | 6.38E-27 | 1.032786 | 0.547 | 0.327 | 1.61E-22 |
| NEXN | 2E-13 | 0.592071 | 0.519 | 0.44 | 5.06E-09 |
| TSC22D1 | 2.49E-11 | 0.700893 | 0.51 | 0.42 | 6.29E-07 |
| ADAMTS1 | 1.6E-08 | 0.855066 | 0.505 | 0.558 | 0.000404 |
| CD302 | 9.79E-14 | 0.598082 | 0.482 | 0.415 | 2.47E-09 |
| TAGLN | 1.36E-15 | 1.628015 | 0.471 | 0.324 | 3.45E-11 |
| MDK | 4.3E-11 | 0.642444 | 0.471 | 0.291 | 1.09E-06 |
| HSPB6 | 4.86E-19 | 0.680471 | 0.465 | 0.352 | 1.23E-14 |
| RBP1 | 2.19E-19 | 0.834128 | 0.451 | 0.209 | 5.55E-15 |
| COL4A2 | 8.03E-24 | 0.940064 | 0.449 | 0.191 | 2.03E-19 |
| GBP1 | 8.72E-08 | 0.789271 | 0.431 | 0.362 | 0.002206 |
| CCL2 | 1.53E-13 | 1.659622 | 0.428 | 0.234 | 3.87E-09 |
| COL18A1 | 1.92E-15 | 0.889494 | 0.425 | 0.286 | 4.84E-11 |
| PHLDA1 | 2.65E-08 | 0.71377 | 0.404 | 0.322 | 0.00067 |
| CYP7B1 | 1.93E-13 | 0.64462 | 0.376 | 0.188 | 4.89E-09 |
| NDRG2 | 2.46E-20 | 0.737922 | 0.369 | 0.143 | 6.22E-16 |
| CYGB | 7.79E-23 | 0.783972 | 0.364 | 0.133 | 1.97E-18 |
| COL4A1 | 1.78E-18 | 0.92709 | 0.359 | 0.138 | 4.5E-14 |
| GAS6 | 1.41E-17 | 0.711539 | 0.355 | 0.168 | 3.56E-13 |
| NR2F1 | 3.07E-21 | 0.860581 | 0.353 | 0.131 | 7.77E-17 |
| PRKAR2B | 8.43E-11 | 0.614476 | 0.35 | 0.204 | 2.13E-06 |
| PLEKHH2 | 1.91E-18 | 0.88314 | 0.345 | 0.158 | 4.82E-14 |
| GUCY1A1 | 1.89E-13 | 0.749041 | 0.344 | 0.204 | 4.79E-09 |
| MYLK | 6.07E-23 | 1.193682 | 0.341 | 0.113 | 1.53E-18 |
| SPON1 | 8.76E-12 | 0.594951 | 0.331 | 0.158 | 2.22E-07 |
| VCAM1 | 6.7E-14 | 0.675541 | 0.325 | 0.166 | 1.7E-09 |
| RAMP1 | 1.53E-22 | 1.155701 | 0.316 | 0.103 | 3.88E-18 |
| CDH11 | 5.64E-13 | 0.624799 | 0.311 | 0.141 | 1.43E-08 |

| | | | | | |
|---|---|---|---|---|---|
| CCL11 | 2.65E-31 | 2.678723 | 0.309 | 0.04 | 6.71E-27 |
| CHCHD10 | 1.71E-16 | 0.612831 | 0.303 | 0.103 | 4.32E-12 |
| KCNN3 | 8.18E-12 | 1.016196 | 0.303 | 0.148 | 2.07E-07 |
| GGT5 | 2.03E-20 | 0.887518 | 0.299 | 0.075 | 5.13E-16 |
| ACTA2 | 3.76E-09 | 1.009069 | 0.297 | 0.226 | 9.5E-05 |
| CTSC | 4.77E-14 | 1.18124 | 0.291 | 0.133 | 1.21E-09 |
| LMOD1 | 1.99E-15 | 0.624765 | 0.281 | 0.098 | 5.03E-11 |
| CCN2 | 6E-11 | 1.064428 | 0.272 | 0.291 | 1.52E-06 |
| ACTN1 | 7.23E-28 | 0.819493 | 0.269 | 0.033 | 1.83E-23 |
| TDO2 | 1.17E-22 | 0.746097 | 0.243 | 0.035 | 2.95E-18 |
| CARMN | 1.64E-19 | 0.737473 | 0.218 | 0.03 | 4.14E-15 |
| FMO2 | 4.46E-17 | 0.862659 | 0.212 | 0.043 | 1.13E-12 |
| FRZB | 1.04E-19 | 0.704692 | 0.201 | 0.023 | 2.63E-15 |
| TNC | 5.38E-17 | 1.00923 | 0.188 | 0.028 | 1.36E-12 |
| COL15A1 | 2.8E-17 | 0.606153 | 0.182 | 0.023 | 7.07E-13 |
| CXCL14 | 6.69E-11 | 2.873001 | 0.176 | 0.07 | 1.69E-06 |
| SCN7A | 2.1E-09 | 0.736626 | 0.168 | 0.058 | 5.3E-05 |
| ADAMDEC1 | 2.2E-15 | 3.185464 | 0.166 | 0.023 | 5.55E-11 |
| IGFBP3 | 2.97E-11 | 0.585254 | 0.135 | 0.176 | 7.52E-07 |
| IGFBP2 | 1.11E-10 | 0.588831 | 0.117 | 0.02 | 2.81E-06 |
| CHI3L1 | 4.08E-11 | 1.524184 | 0.107 | 0.01 | 1.03E-06 |
| CCL19 | 7.72E-11 | 1.652495 | 0.068 | 0 | 1.95E-06 |

**Supplementary Table 2. Gene markers that define fibroblast cell clusters C9 and C12 derived from either stricturing Crohns disease (SCD) or non-stricturing CD (NSCD).** List of all differentially expressed genes identified using the findmarkers function (MAST algorithm) to compare cluster (C)9 SCD, C9 NSCD, C12 SCD and C12 NSCD fibroblast populations. Each table has been ordered on pct.1 and pct.2 columns.

**SCD C9 compared to SCD C12**

| Gene | p_val | avg_log2FC | pct.1 | pct.2 | p_val_adj |
|---|---|---|---|---|---|
| JUN | 6.23E-19 | 0.643443 | 0.948 | 0.957 | 1.58E-14 |
| FOSB | 3.51E-20 | 0.694453 | 0.917 | 0.766 | 8.88E-16 |
| CALD1 | 4.25E-43 | 0.729076 | 0.889 | 0.954 | 1.07E-38 |
| CXCL12 | 6.72E-14 | 0.593581 | 0.861 | 0.828 | 1.7E-09 |
| MGP | 1.11E-23 | 0.590613 | 0.846 | 0.976 | 2.81E-19 |
| IGFBP4 | 6.21E-21 | 0.682621 | 0.83 | 0.898 | 1.57E-16 |
| SAT1 | 2.88E-08 | 0.70402 | 0.75 | 0.728 | 0.000727 |
| PTGDS | 6.77E-42 | 2.068807 | 0.748 | 0.439 | 1.71E-37 |
| A2M | 1.11E-24 | 1.067084 | 0.733 | 0.542 | 2.81E-20 |
| APOE | 1E-13 | 1.005386 | 0.722 | 0.577 | 2.53E-09 |
| IGKC | 5.61E-27 | 5.579525 | 0.693 | 0.392 | 1.42E-22 |
| BTG1 | 7.19E-14 | 0.73884 | 0.693 | 0.614 | 1.82E-09 |
| TPM4 | 4.96E-12 | 0.61689 | 0.687 | 0.697 | 1.25E-07 |
| TPM2 | 1.45E-44 | 1.377476 | 0.685 | 0.448 | 3.65E-40 |
| C7 | 1.28E-19 | 1.058961 | 0.659 | 0.382 | 3.23E-15 |
| IGF1 | 5.57E-12 | 0.69066 | 0.637 | 0.549 | 1.41E-07 |
| EMILIN1 | 2.58E-33 | 1.074964 | 0.602 | 0.353 | 6.52E-29 |
| GREM1 | 6.57E-12 | 1.038143 | 0.585 | 0.398 | 1.66E-07 |
| NEXN | 4.98E-18 | 0.698882 | 0.561 | 0.432 | 1.26E-13 |
| IGHA1 | 6.47E-16 | 4.749849 | 0.557 | 0.343 | 1.64E-11 |
| TSC22D1 | 8.67E-11 | 0.661108 | 0.546 | 0.42 | 2.19E-06 |
| PALLD | 2.48E-12 | 0.606316 | 0.543 | 0.503 | 6.27E-08 |
| MDK | 2.39E-19 | 0.808437 | 0.537 | 0.296 | 6.03E-15 |
| TAGLN | 5.12E-26 | 1.795757 | 0.515 | 0.336 | 1.29E-21 |
| HSPB6 | 1.35E-15 | 0.645346 | 0.498 | 0.361 | 3.42E-11 |
| RBP1 | 2E-18 | 0.761547 | 0.491 | 0.253 | 5.07E-14 |
| COL18A1 | 1.69E-19 | 0.959891 | 0.476 | 0.289 | 4.27E-15 |
| GBP1 | 2.13E-07 | 0.752597 | 0.465 | 0.356 | 0.005395 |
| MYLK | 2.54E-35 | 1.376355 | 0.426 | 0.117 | 6.43E-31 |
| RAMP1 | 3.73E-38 | 1.394536 | 0.407 | 0.098 | 9.44E-34 |
| IGLC2 | 5.15E-10 | 1.656164 | 0.404 | 0.236 | 1.3E-05 |
| NDRG2 | 2.44E-14 | 0.604347 | 0.398 | 0.191 | 6.16E-10 |
| CTGF | 5.74E-13 | 1.204431 | 0.385 | 0.201 | 1.45E-08 |
| NR2F1 | 2.71E-17 | 0.772674 | 0.376 | 0.182 | 6.85E-13 |
| CCL11 | 1.48E-31 | 2.466773 | 0.37 | 0.077 | 3.73E-27 |
| PLEKHH2 | 3.53E-11 | 0.614261 | 0.367 | 0.2 | 8.93E-07 |
| CYR61 | 4.92E-10 | 0.865362 | 0.363 | 0.188 | 1.24E-05 |

| | | | | | |
|---|---|---|---|---|---|
| GGT5 | 8.19E-22 | 0.814276 | 0.35 | 0.105 | 2.07E-17 |
| TSPYL2 | 1.31E-10 | 0.616939 | 0.343 | 0.182 | 3.31E-06 |
| CDH11 | 8.48E-12 | 0.600374 | 0.339 | 0.172 | 2.14E-07 |
| ACTA2 | 1.03E-09 | 0.810919 | 0.333 | 0.22 | 2.6E-05 |
| ACTN1 | 2.35E-28 | 0.858359 | 0.322 | 0.065 | 5.95E-24 |
| LMOD1 | 1.7E-14 | 0.615023 | 0.315 | 0.129 | 4.29E-10 |
| KCNN3 | 8.21E-08 | 0.810225 | 0.315 | 0.188 | 0.002076 |
| CNN2 | 2.37E-21 | 0.679976 | 0.287 | 0.072 | 6E-17 |
| CARMN | 3.21E-20 | 0.586355 | 0.261 | 0.055 | 8.12E-16 |
| FRZB | 6.34E-25 | 0.81541 | 0.252 | 0.038 | 1.6E-20 |
| CCN2 | 2.14E-20 | 1.124219 | 0.243 | 0.308 | 5.4E-16 |
| TNC | 5.74E-10 | 0.648749 | 0.202 | 0.067 | 1.45E-05 |
| JCHAIN | 6.75E-09 | 4.8262 | 0.198 | 0.1 | 0.000171 |
| IGHA2 | 6.29E-08 | 2.201564 | 0.17 | 0.095 | 0.001591 |
| CHI3L1 | 4.13E-20 | 1.874169 | 0.148 | 0.009 | 1.04E-15 |
| IGFBP2 | 7.68E-12 | 0.653282 | 0.143 | 0.029 | 1.94E-07 |

**SCD C12 compared to SCD C9**

| Gene | p_val | avg_log2FC | pct.1 | pct.2 | p_val_adj |
|---|---|---|---|---|---|
| CST3 | 1.4E-24 | 0.596117 | 1 | 0.937 | 3.55E-20 |
| PLAC9 | 3.96E-51 | 1.172868 | 0.996 | 0.71 | 1E-46 |
| C1R | 8.17E-24 | 0.648557 | 0.996 | 0.933 | 2.07E-19 |
| CFD | 1.91E-23 | 0.79448 | 0.996 | 0.887 | 4.82E-19 |
| IGFBP6 | 1.16E-46 | 1.324019 | 0.991 | 0.799 | 2.94E-42 |
| FBN1 | 5.8E-39 | 1.224798 | 0.991 | 0.833 | 1.47E-34 |
| S100A10 | 9.9E-34 | 0.824102 | 0.991 | 0.807 | 2.5E-29 |
| S100A4 | 8.1E-32 | 0.814795 | 0.991 | 0.849 | 2.05E-27 |
| S100A6 | 2.46E-23 | 0.662737 | 0.991 | 0.949 | 6.21E-19 |
| TMSB10 | 1.86E-20 | 0.635858 | 0.991 | 0.954 | 4.69E-16 |
| TIMP1 | 1.64E-30 | 0.959598 | 0.987 | 0.881 | 4.16E-26 |
| GSN | 2E-17 | 0.647742 | 0.987 | 0.907 | 5.05E-13 |
| TNXB | 8.97E-42 | 1.197756 | 0.983 | 0.808 | 2.27E-37 |
| FSTL1 | 2.29E-24 | 0.856965 | 0.979 | 0.789 | 5.8E-20 |
| LTBP4 | 1.52E-22 | 0.720081 | 0.966 | 0.812 | 3.86E-18 |
| TIMP3 | 1.27E-21 | 0.702551 | 0.966 | 0.767 | 3.22E-17 |
| C3 | 1.37E-10 | 0.6075 | 0.962 | 0.861 | 3.45E-06 |
| ANXA2 | 3.22E-25 | 0.744397 | 0.957 | 0.819 | 8.14E-21 |
| EFEMP1 | 4.25E-22 | 0.688977 | 0.957 | 0.748 | 1.08E-17 |
| SFRP2 | 7.55E-14 | 0.714755 | 0.944 | 0.772 | 1.91E-09 |
| CD248 | 5.63E-46 | 1.228269 | 0.94 | 0.576 | 1.42E-41 |
| S100A11 | 2.14E-16 | 0.610541 | 0.932 | 0.788 | 5.42E-12 |
| MFAP5 | 9.79E-48 | 1.182989 | 0.927 | 0.44 | 2.48E-43 |
| EMP3 | 4.9E-19 | 0.80073 | 0.902 | 0.713 | 1.24E-14 |
| PCOLCE | 7.69E-16 | 0.64405 | 0.897 | 0.709 | 1.94E-11 |
| GPX3 | 4.15E-14 | 0.603599 | 0.897 | 0.691 | 1.05E-09 |

| | | | | | |
|---|---|---|---|---|---|
| FN1 | 1.53E-27 | 0.983479 | 0.88 | 0.53 | 3.88E-23 |
| SEMA3C | 7.2E-56 | 1.470149 | 0.876 | 0.33 | 1.82E-51 |
| FBLN2 | 4.26E-27 | 0.816174 | 0.876 | 0.503 | 1.08E-22 |
| CD55 | 6.56E-37 | 1.143964 | 0.868 | 0.452 | 1.66E-32 |
| SLIT3 | 1.91E-18 | 0.657425 | 0.863 | 0.612 | 4.84E-14 |
| CLU | 5.35E-22 | 0.907026 | 0.859 | 0.572 | 1.35E-17 |
| SH3BGRL3 | 6.27E-28 | 1.023911 | 0.838 | 0.584 | 1.59E-23 |
| MT2A | 2.31E-08 | 0.994664 | 0.833 | 0.672 | 0.000583 |
| ISLR | 4.91E-16 | 0.691261 | 0.829 | 0.611 | 1.24E-11 |
| PLA2G2A | 4.95E-27 | 1.395943 | 0.812 | 0.447 | 1.25E-22 |
| SCARA5 | 6.79E-36 | 1.208394 | 0.795 | 0.352 | 1.72E-31 |
| PIGT | 3.8E-23 | 0.767854 | 0.786 | 0.466 | 9.61E-19 |
| HTRA3 | 2.25E-18 | 1.03424 | 0.765 | 0.487 | 5.68E-14 |
| ADAMTS5 | 1.66E-32 | 1.322687 | 0.761 | 0.344 | 4.21E-28 |
| CADM3 | 8.21E-30 | 0.985186 | 0.756 | 0.344 | 2.08E-25 |
| CLTB | 2.43E-19 | 0.747508 | 0.756 | 0.48 | 6.13E-15 |
| UAP1 | 1.68E-20 | 0.886572 | 0.752 | 0.424 | 4.24E-16 |
| GAS7 | 7.32E-22 | 0.754554 | 0.748 | 0.409 | 1.85E-17 |
| ABLIM1 | 2.63E-21 | 0.764961 | 0.744 | 0.395 | 6.66E-17 |
| CLEC3B | 2.96E-37 | 1.155105 | 0.709 | 0.248 | 7.5E-33 |
| SERPINE2 | 6.26E-09 | 0.654068 | 0.684 | 0.458 | 0.000158 |
| ZNF385A | 2.6E-24 | 0.797351 | 0.679 | 0.317 | 6.57E-20 |
| PTGIS | 1.04E-13 | 0.696719 | 0.675 | 0.404 | 2.63E-09 |
| METRNL | 3.69E-21 | 0.808774 | 0.662 | 0.341 | 9.34E-17 |
| TPPP3 | 8.58E-17 | 0.80756 | 0.662 | 0.354 | 2.17E-12 |
| DBN1 | 1.37E-25 | 0.900308 | 0.654 | 0.284 | 3.46E-21 |
| OSR2 | 2.53E-14 | 0.682238 | 0.65 | 0.366 | 6.4E-10 |
| ACKR3 | 9.46E-17 | 0.828336 | 0.628 | 0.317 | 2.39E-12 |
| PRSS23 | 1E-21 | 0.801049 | 0.607 | 0.258 | 2.53E-17 |
| SLPI | 3.39E-25 | 1.396333 | 0.603 | 0.247 | 8.58E-21 |
| ADAMTSL4 | 2.02E-18 | 0.601411 | 0.543 | 0.228 | 5.1E-14 |
| PI16 | 6.13E-20 | 0.912871 | 0.534 | 0.208 | 1.55E-15 |
| SEMA3E | 3.31E-18 | 0.814392 | 0.491 | 0.188 | 8.37E-14 |
| EMILIN2 | 2.77E-14 | 0.598625 | 0.479 | 0.212 | 7E-10 |
| AC245595.1 | 1.16E-14 | 0.658685 | 0.457 | 0.209 | 2.94E-10 |
| PCOLCE2 | 5.31E-12 | 0.944578 | 0.457 | 0.216 | 1.34E-07 |
| ITGA11 | 1.37E-11 | 0.617691 | 0.444 | 0.212 | 3.46E-07 |
| GALNT15 | 4.83E-12 | 0.670651 | 0.342 | 0.136 | 1.22E-07 |
| WISP2 | 2.2E-11 | 0.664695 | 0.321 | 0.116 | 5.56E-07 |
| KRT24 | 1.89E-24 | 0.703316 | 0.244 | 0.021 | 4.79E-20 |

**NSCD Cluster 9 compared to NSCD 12**

| Gene | p_val | avg_log2FC | pct.1 | pct.2 | p_val_adj |
|---|---|---|---|---|---|
| VIM | 2.38E-15 | 0.661329 | 0.995 | 0.976 | 6.01E-11 |
| RPL13A1 | 2.7E-19 | 0.653872 | 0.967 | 0.958 | 6.83E-15 |
| SPARCL1 | 2.12E-16 | 0.797342 | 0.951 | 0.889 | 5.36E-12 |
| JUNB | 1.72E-15 | 0.814196 | 0.923 | 0.869 | 4.34E-11 |
| CEBPD | 5.28E-15 | 0.742794 | 0.923 | 0.821 | 1.34E-10 |
| RPS291 | 4.74E-16 | 0.631995 | 0.918 | 0.861 | 1.2E-11 |
| RPL23A1 | 1.95E-18 | 0.716642 | 0.907 | 0.846 | 4.93E-14 |
| RPL381 | 4.97E-16 | 0.652227 | 0.874 | 0.78 | 1.26E-11 |
| RPL311 | 8.2E-13 | 0.620199 | 0.825 | 0.763 | 2.07E-08 |
| IGFBP7 | 1.82E-12 | 0.774516 | 0.82 | 0.691 | 4.6E-08 |
| RPL171 | 1.58E-15 | 0.94199 | 0.809 | 0.601 | 4E-11 |
| PLCG21 | 2.62E-21 | 1.036693 | 0.803 | 0.681 | 6.62E-17 |
| RPL36A | 8.27E-11 | 0.588824 | 0.803 | 0.775 | 2.09E-06 |
| ABCA8 | 7.17E-12 | 0.759038 | 0.705 | 0.521 | 1.81E-07 |
| C71 | 1.47E-09 | 0.805483 | 0.694 | 0.464 | 3.71E-05 |
| SOD21 | 1.98E-08 | 0.801065 | 0.694 | 0.69 | 0.0005 |
| GABARAP1 | 4.81E-10 | 0.626639 | 0.689 | 0.45 | 1.22E-05 |
| EMP1 | 5.47E-10 | 0.758919 | 0.689 | 0.52 | 1.38E-05 |
| ANGPTL1 | 2.83E-08 | 0.595601 | 0.65 | 0.519 | 0.000717 |
| CD302 | 3.23E-13 | 0.818508 | 0.639 | 0.417 | 8.18E-09 |
| PTN1 | 1.79E-08 | 0.650815 | 0.623 | 0.466 | 0.000454 |
| RNASE41 | 3.83E-09 | 0.602899 | 0.607 | 0.385 | 9.68E-05 |
| GAS5 | 8.39E-17 | 0.883573 | 0.574 | 0.273 | 2.12E-12 |
| SNHG61 | 4.03E-12 | 0.79237 | 0.525 | 0.273 | 1.02E-07 |
| ABCA9 | 1.06E-11 | 0.849184 | 0.497 | 0.329 | 2.68E-07 |
| FABP6 | 4.64E-09 | 1.049292 | 0.492 | 0.29 | 0.000117 |
| AL627171.21 | 6.77E-11 | 0.71726 | 0.448 | 0.217 | 1.71E-06 |
| ASPN | 2.07E-07 | 0.605787 | 0.443 | 0.239 | 0.005244 |
| GAS6 | 5.4E-09 | 0.613279 | 0.415 | 0.255 | 0.000137 |
| ID3 | 3.93E-07 | 0.91891 | 0.383 | 0.23 | 0.009943 |
| APOA1 | 2.09E-12 | 0.781535 | 0.344 | 0.131 | 5.3E-08 |
| FMO2 | 3.33E-12 | 0.957426 | 0.311 | 0.112 | 8.42E-08 |
| IGFBP3 | 1.36E-10 | 1.353383 | 0.262 | 0.127 | 3.45E-06 |
| FABP4 | 1.44E-08 | 0.710746 | 0.115 | 0.015 | 0.000363 |

**NSCD C12 compared to NSCD C9**

| Gene | p_val | avg_log2FC | pct.1 | pct.2 | p_val_adj |
|---|---|---|---|---|---|
| MFAP51 | 1.09E-75 | 2.009796 | 1 | 0.465 | 2.75E-71 |
| S100A101 | 5.05E-47 | 1.267946 | 1 | 0.82 | 1.28E-42 |
| IGFBP61 | 7.63E-41 | 1.195882 | 1 | 0.813 | 1.93E-36 |
| TMSB4X | 2.23E-36 | 0.976345 | 1 | 0.97 | 5.65E-32 |
| CFD1 | 7.39E-19 | 0.737359 | 1 | 0.895 | 1.87E-14 |

| | | | | | |
|---|---|---|---|---|---|
| B2M | 1.71E-17 | 0.601636 | 1 | 0.989 | 4.32E-13 |
| FBN11 | 4.98E-44 | 1.379843 | 0.994 | 0.845 | 1.26E-39 |
| PLAC91 | 4.04E-37 | 1.07568 | 0.994 | 0.733 | 1.02E-32 |
| S100A61 | 1.04E-24 | 0.807697 | 0.994 | 0.952 | 2.62E-20 |
| TNXB1 | 1.43E-20 | 0.629549 | 0.994 | 0.82 | 3.61E-16 |
| TIMP31 | 1.19E-28 | 0.928094 | 0.988 | 0.779 | 3.02E-24 |
| MFAP4 | 1.2E-20 | 0.793312 | 0.988 | 0.829 | 3.04E-16 |
| S100A41 | 1.41E-28 | 0.970222 | 0.982 | 0.862 | 3.56E-24 |
| FN11 | 3.54E-46 | 1.593593 | 0.976 | 0.54 | 8.95E-42 |
| S100A111 | 1.36E-15 | 0.659436 | 0.97 | 0.792 | 3.44E-11 |
| IGFBP51 | 1.38E-09 | 0.696902 | 0.97 | 0.818 | 3.49E-05 |
| FBLN21 | 1.12E-42 | 1.178326 | 0.963 | 0.517 | 2.84E-38 |
| CD2481 | 2.77E-27 | 0.722271 | 0.963 | 0.601 | 7.01E-23 |
| FSTL11 | 3.7E-31 | 1.108986 | 0.957 | 0.808 | 9.35E-27 |
| ANXA1 | 5.17E-14 | 0.58978 | 0.957 | 0.741 | 1.31E-09 |
| EMP31 | 1.04E-23 | 0.836194 | 0.945 | 0.719 | 2.62E-19 |
| EFEMP11 | 4.13E-16 | 0.647214 | 0.945 | 0.767 | 1.04E-11 |
| CD991 | 1.18E-19 | 0.68672 | 0.939 | 0.79 | 2.99E-15 |
| SCARA51 | 3.22E-45 | 1.091344 | 0.927 | 0.363 | 8.15E-41 |
| CD551 | 2.27E-45 | 1.712277 | 0.921 | 0.475 | 5.73E-41 |
| SEMA3C1 | 1.28E-44 | 1.328471 | 0.921 | 0.365 | 3.24E-40 |
| SDC21 | 8.89E-29 | 0.919619 | 0.921 | 0.559 | 2.25E-24 |
| TXNIP | 2.44E-11 | 0.693703 | 0.921 | 0.791 | 6.16E-07 |
| CLIC1 | 2.05E-18 | 0.693177 | 0.909 | 0.636 | 5.18E-14 |
| UAP11 | 3.64E-36 | 1.176282 | 0.89 | 0.424 | 9.22E-32 |
| KLF4 | 7.73E-18 | 0.593983 | 0.89 | 0.561 | 1.96E-13 |
| PPIC1 | 1.69E-29 | 1.003013 | 0.872 | 0.519 | 4.28E-25 |
| KLF2 | 2.53E-18 | 0.980407 | 0.872 | 0.595 | 6.39E-14 |
| ACKR31 | 6.11E-43 | 1.235495 | 0.86 | 0.299 | 1.54E-38 |
| TXN | 3.28E-16 | 0.608984 | 0.86 | 0.558 | 8.31E-12 |
| SH3BGRL31 | 2.57E-14 | 0.655053 | 0.86 | 0.6 | 6.5E-10 |
| C17orf581 | 4.71E-49 | 1.749344 | 0.854 | 0.352 | 1.19E-44 |
| TPPP31 | 4.62E-36 | 1.262775 | 0.835 | 0.347 | 1.17E-31 |
| PLA2G2A1 | 3.78E-18 | 0.786758 | 0.835 | 0.472 | 9.56E-14 |
| CLEC3B1 | 2.83E-43 | 1.178453 | 0.829 | 0.262 | 7.16E-39 |
| ITM2A1 | 3.19E-30 | 1.08171 | 0.817 | 0.368 | 8.07E-26 |
| CREB51 | 1.24E-26 | 0.998728 | 0.811 | 0.382 | 3.14E-22 |
| PLCG2 | 6.22E-10 | 0.669062 | 0.805 | 0.683 | 1.57E-05 |
| CRIP11 | 1.53E-29 | 1.39087 | 0.799 | 0.363 | 3.88E-25 |
| EBF11 | 1.86E-15 | 0.626482 | 0.799 | 0.469 | 4.7E-11 |
| ADAMTS51 | 1.85E-25 | 1.133022 | 0.793 | 0.372 | 4.68E-21 |
| CD341 | 1.02E-18 | 0.723396 | 0.793 | 0.433 | 2.57E-14 |
| REXO2 | 3.66E-15 | 0.689125 | 0.793 | 0.547 | 9.25E-11 |
| METRNL1 | 1.44E-26 | 0.912153 | 0.787 | 0.343 | 3.64E-22 |
| CADM31 | 5.79E-24 | 0.897351 | 0.787 | 0.372 | 1.46E-19 |
| PCOLCE21 | 3.2E-51 | 1.386683 | 0.78 | 0.174 | 8.09E-47 |
| MEDAG1 | 3.62E-15 | 0.742388 | 0.774 | 0.45 | 9.16E-11 |
| UGP2 | 2.41E-19 | 0.901331 | 0.768 | 0.432 | 6.09E-15 |
| ADI11 | 1.44E-13 | 0.656622 | 0.744 | 0.448 | 3.64E-09 |
| DBN11 | 1.14E-24 | 0.781909 | 0.732 | 0.299 | 2.88E-20 |

| | | | | | |
|---|---|---|---|---|---|
| ARHGAP291 | 8.6E-22 | 0.881614 | 0.72 | 0.336 | 2.18E-17 |
| EEF1G | 1.11E-12 | 0.742158 | 0.713 | 0.413 | 2.81E-08 |
| GAS71 | 6.01E-12 | 0.69005 | 0.713 | 0.442 | 1.52E-07 |
| F101 | 4.15E-13 | 0.613399 | 0.707 | 0.396 | 1.05E-08 |
| YWHAH1 | 1.38E-13 | 0.689604 | 0.695 | 0.388 | 3.49E-09 |
| SERPINE21 | 2.6E-07 | 0.597653 | 0.689 | 0.475 | 0.006581 |
| PI161 | 1.78E-29 | 1.45941 | 0.671 | 0.209 | 4.49E-25 |
| LRRC171 | 1.34E-26 | 0.966936 | 0.671 | 0.23 | 3.39E-22 |
| ZNF385A1 | 3.61E-13 | 0.622091 | 0.665 | 0.349 | 9.13E-09 |
| PRSS231 | 5.59E-21 | 0.920485 | 0.659 | 0.276 | 1.41E-16 |
| SEMA3E1 | 2.59E-30 | 1.041685 | 0.646 | 0.184 | 6.54E-26 |
| UGDH1 | 2.33E-19 | 0.742989 | 0.646 | 0.273 | 5.89E-15 |
| APBB1IP | 1.98E-19 | 0.632444 | 0.64 | 0.26 | 5.01E-15 |
| TBC1D12 | 5.17E-22 | 0.775019 | 0.628 | 0.233 | 1.31E-17 |
| VASN1 | 3.42E-20 | 0.768954 | 0.628 | 0.246 | 8.66E-16 |
| PROCR1 | 8.54E-29 | 1.090049 | 0.622 | 0.18 | 2.16E-24 |
| LINC011331 | 3.5E-24 | 0.876279 | 0.616 | 0.204 | 8.85E-20 |
| EMILIN21 | 2.56E-21 | 0.606947 | 0.598 | 0.211 | 6.47E-17 |
| LOXL11 | 1.07E-15 | 0.867468 | 0.598 | 0.294 | 2.72E-11 |
| SLPI1 | 5.36E-13 | 0.792674 | 0.585 | 0.278 | 1.35E-08 |
| GFPT21 | 1.23E-14 | 0.688395 | 0.579 | 0.255 | 3.11E-10 |
| VEGFA | 2.85E-11 | 0.706343 | 0.573 | 0.293 | 7.21E-07 |
| C12orf751 | 4.33E-21 | 0.600283 | 0.537 | 0.166 | 1.09E-16 |
| LINC02802 | 8.75E-18 | 0.787962 | 0.537 | 0.193 | 2.21E-13 |
| TLE5 | 1.82E-12 | 0.613623 | 0.53 | 0.236 | 4.62E-08 |
| MICOS10 | 4.02E-12 | 0.591859 | 0.524 | 0.235 | 1.02E-07 |
| SHISA3 | 6.92E-14 | 0.712518 | 0.512 | 0.21 | 1.75E-09 |
| TRIO1 | 1.08E-11 | 0.682592 | 0.512 | 0.242 | 2.74E-07 |
| LEPR | 8.7E-17 | 0.612609 | 0.494 | 0.169 | 2.2E-12 |
| NTM | 1.79E-19 | 0.656667 | 0.445 | 0.117 | 4.52E-15 |
| CPE | 1.11E-07 | 0.786848 | 0.433 | 0.242 | 0.002797 |
| DBNDD2 | 1.16E-16 | 0.667717 | 0.366 | 0.09 | 2.92E-12 |
| CD24 | 6.72E-12 | 0.894469 | 0.305 | 0.087 | 1.7E-07 |
| H19 | 1.5E-13 | 1.283641 | 0.268 | 0.064 | 3.79E-09 |
| HAS1 | 1.16E-08 | 1.121996 | 0.207 | 0.068 | 0.000293 |
| HBEGF | 8.98E-10 | 0.597739 | 0.189 | 0.038 | 2.27E-05 |
